# Low potential electron generating enzymes serve different functions during aerobic nitrogen fixation in *Azotobacter vinelandii*

**DOI:** 10.1101/2021.10.06.463449

**Authors:** Alexander B. Alleman, Florence Mus, John W. Peters

**Affiliations:** Institute of Biological Chemistry, Washington State University, Pullman, Washington, USA

## Abstract

Biological nitrogen fixation requires large amounts of energy in the form of ATP and low potential electrons to overcome the high activation barrier for cleavage of the dinitrogen triple bond. The model aerobic nitrogen-fixing bacteria, *Azotobacter vinelandii,* generates low potential electrons in the form of reduced ferredoxin (Fd) and flavodoxin (Fld) using two distinct mechanisms via the enzyme complexes Rnf and Fix. Both Rnf and Fix are expressed during nitrogen fixation, and deleting either *rnf1* or *fix* genes has little effect on diazotrophic growth. However, deleting both *rnf1* and *fix* eliminates the ability to grow diazotrophically. Rnf and Fix both use NADH as a source of electrons, but overcoming the energetics of NADH’s endergonic reduction of Fd/Fld is accomplished through different mechanisms. Rnf harnesses free energy from the proton motive force, whereas Fix uses electron bifurcation to effectively couple the endergonic reduction of Fd/Fld to the exergonic reduction of quinone. Different stoichiometries and gene expression analyses indicate specific roles for the two reactions under different conditions. In this work, complementary physiological studies and thermodynamic modeling reveal how Rnf and Fix simultaneously balance redox homeostasis in various conditions. Specifically, the Fix complex is required for efficient growth under low oxygen concentrations, while Rnf sustains homeostasis and delivers sufficient reduced Fd to nitrogenase under standard conditions. This work provides a framework for understanding how the production of low potential electrons sustains robust nitrogen fixation in various conditions.

**Importance:** The availability of fixed nitrogen is critical for life in many ecosystems, from hot springs to agriculture. Due to the energy demands of biological nitrogen fixation, organisms must tailor their metabolism during diazotrophic growth to deliver energy requirements to nitrogenase in the form of ATP and low potential electrons. Therefore, a complete understanding of diazotrophic energy metabolism and redox homeostasis is required to understand the impact on ecological communities or to promote crop growth in agriculture through engineered diazotrophs.

## Introduction

Organisms that fix nitrogen, termed diazotrophs, do so through the activity of the enzyme nitrogenase, which catalyzes the reduction of atmospheric dinitrogen to ammonia in a complex reaction that uses large amounts of energy in the form of ATP and low potential electrons (Eq 1) (1).

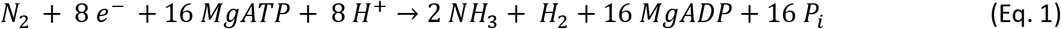

Nitrogenase requires electrons at a mid-point potential of at least −400 mV (2). This is especially a problem for diazotrophic bacteria that fix nitrogen with aerobic metabolisms where most electrons are exchanged as NAD^+^/NADH couples (3). Because electrons from NADH cannot reduce nitrogenase as the mid-point potential of NADH (−320 mV) is too high. Thus, small redox proteins flavodoxin (Fld) and ferredoxin (Fd), with a mid-point potential ~-450-500 mV, serve as electron donors to nitrogenase *in vivo* (4–8). In aerobic metabolisms, the endergonic reduction of Fd/Fld from NADH must be coupled to an exergonic reaction to maintain nitrogen fixation.

*Azotobacter vinelandii* is one of the most robust diazotrophs due to its ability to fix nitrogen in various oxygen concentrations even though nitrogenase is extremely sensitive to oxygen (9, 10). The capacity to grow in various conditions is partly due to *A. vinelandii’s* complex electron chain consisting of multiple branches. These branches allow *A. vinelandii* to partition electrons from NADH down three paths; 1) a fully proton-coupled electron transport chain (ETC) to maximize ATP production, 2) a partially proton-coupled ETC to maximize O_2_ consumption and protect nitrogenase, or 3) production of low potential electrons Fd/Fld. While roles of the branched ETC and terminal oxidases are beginning to be understood, little is known about the partitioning of electrons to reduced Fd/Fld.

Production of reduced Fd/Fld in *A. vinelandii* is catalyzed by two enzyme complexes called Rnf1 and Fix. Both complexes oxidize NADH and reduce Fd/Fld by using distinct mechanisms. Rnf1, originally named from <u>R</u>hodobacter <u>N</u>itrogen <u>F</u>ixation, couples the endergonic transfer of electrons from NADH to Fd/Fld with the exergonic transport of ions/protons by the proton motive force (*pmf*) (11, 12). Rnfs have been identified in many bacteria; in *Acetobacterium woodii,* the reduction of NAD^+^ to NADH from Fd/Fld is coupled to the pumping of Na^+^ ions (13, 14). In *A. vinelandii* and *Rhodobacter capsulatus,* overexpression of *rnf* genes increases *in vivo* nitrogenase activity (12, 15). *A. vinelandii* contains two *rnf* clusters that are differentially regulated. The *rnf1* genes are within the nitrogen fixation (*nif*) gene cluster and are up-regulated under nitrogen-fixing conditions, while *rnf2* genes are constitutively expressed and are not affected by the nitrogen status (16, 17). Relative to wild-type (WT), *rnf1* deletion mutants exhibit longer growth lags during the transition to diazotrophic growth; overall growth rates are comparable (16). *A. vinelandii* also generates Fd/Fld during nitrogen fixation through the enzyme complex Fix, which is present in rhizobia and other free-living diazotrophs (18–20). The Fix complex overcomes the energy barrier of Fd/Fld reduction by using flavin-based electron bifurcation, which couples the exergonic reduction of quinone with the endergonic reduction of Fd/Fld (21–23).

In *A. vinelandii*, deletion of either *rnf1* or *fix* has a limited effect on diazotrophic growth in standard laboratory conditions. However, deleting both *rnf1* and *fix* eliminates the mutant’s ability to grow diazotrophically (22). This implies that both Rnf1 and Fix are used to transport electrons to nitrogenase through Fd/Fld and are somewhat redundant in function, therefore, compensating for each other under laboratory growth conditions. Even though the enzymes appear redundant, the energetics of the endergonic reduction of Fd/Fld by NADH is accomplished through different mechanisms. Rnf uses the free energy from the *pmf* and Fix couples the reduction of Fd/Fld to quinone. Gene expression analysis indicates different regulatory patterns for each enzyme complex. The *fixABCX* genes are up-regulated under alternative nitrogenase conditions, expressed when Mo or Mo and V are limited in the environment (17). While *rnfABCDEG1* genes are up-regulated when regulation of nitrogenase is interrupted, causing an ammonia excretion phenotype (15, 24). Here we demonstrate distinct roles for Rnf and Fix under different oxygen concentrations and metal availability. Growth rates and physiological parameters were coupled with a simple thermodynamic model to propose how these two enzymes balance the electron transfer to nitrogenase while maintaining ATP production.

## Results and discussion

In previous work, strains Δ*rnf1* and Δ*fix* showed little difference in their ability to grow diazotrophically relative to WT (22). While this shows functional redundancy, the favored mechanism and/or individual roles for Rnf and Fix in nitrogen fixation remained unknown. Under standard laboratory conditions in media in which carbon is not limiting, the growth of batch cultures is limited by the nitrogen fixation rate. Under these conditions, added ammonia increases the growth rate, and as such, effects on nitrogen fixation rate should be manifested in changes in the growth rate. However, once carbon or oxygen concentrations become limited, *A. vinelandii* can no longer maximize nitrogen fixation. *A. vinelandii’s* ability to maintain electron transport to nitrogenase in various conditions would be evolutionarily advantageous. Under standard conditions of carbon and oxygen excess at a rotation rate of 200 RPM, there is no difference in growth rate between Δ*rnf1*, Δ*fix,* or WT under Mo-dependent diazotrophic conditions (Table 1). However, when the aeration rate is lowered to 90 RPM, there is a dramatic growth decrease for WT as the growth shifts from nitrogenase-limited to oxygen-limited conditions. The Δ*rnf1* strain grown at 90 RPM sees a similar reduction in growth rate as WT. The Δ*fix* strain grown under low aeration diazotrophic conditions was even more profoundly impacted, having a 3-4 times slower growth rate compared to Δ*rnf1* or WT (Table 1). The observed growth phenotypes were not related to shaking or mixing rate as they can be reproduced in 6-well plates under constant shaking but a low oxygen atmosphere (Table S1). These observations indicate a specific role for Fix under oxygen-limited conditions. In low oxygen, when only Rnf is present (Δ*fix* strain), electron transport to nitrogenase was limiting. In contrast, similar growth rates at high aeration mean both Rnf and Fix can maintain electron transport under oxygen and carbon excess conditions. When *A. vinelandii* is grown in oxygen-limited conditions, a decrease in respiration increases NADH and NADPH concentrations, thus inhibiting citrate synthase and isocitrate dehydrogenase and increasing the concentration of acetyl-CoA available for poly(3-hydroxybutyrate) (P(3HB)) biosynthesis (25–27). The increase of P(3HB) and decrease in respiration improve the biomass yield of *A. vinelandii,* making low oxygen conditions ideal for biopolymer production (28–30). The oxygen concentration affects the metabolic fluxes through carbon metabolism and influences the NAD(P)H/NAD(P)^+^ ratios (26). When the cells become oxygen-limited, the increased NADH would favor the Fix mechanism, while a decrease of respiration would hinder the use of *pmf* by Rnf.

**Table 1.** Generation times for WT, Δ*rnf1 and* Δ*fix* strains under Mo-dependent diazotrophic growth in high and low aeration conditions. Cells were grown in 200 mL of B or BN medium in a 500 mL baffled Erlenmeyer flask either at 200 RPM for high aeration or 90 RPM for low aeration.

| | | WT | $\Delta rnf1$ | $\Delta fix$ |
| --- | --- | --- | --- | --- |
|  |  | Generation Time (hrs) |  |  |
| Diazotrophic | High Aeration | $3.1 \pm 0.1$ | $3.1 \pm 0.2$ | $3.2 \pm 0.1$ |
| | Low Aeration | $12.3 \pm 1.9$ | $11.6 \pm 0.8$ | $36.6 \pm 13.2$ |
| Ammonia supplemented | High Aeration | $2.6 \pm 0.2$ | $3.0 \pm 0.1$ | $2.4 \pm 0.3$ |
| | Low Aeration | $9.9 \pm 2.1$ | $10.4 \pm 1.2$ | $12.5 \pm 3.6$ |

To further understand the dynamics of both Rnf and Fix under higher oxygen concentrations, physiological parameters were measured. Significant energy decoupling is required to maintain nitrogen fixation in high oxygen and carbon conditions (31–33). Therefore, the ability to simultaneously maximize Fd/Fld and oxygen reduction would be advantageous. To measure Fd/Fld reduction, a metronidazole susceptibility assay was used as a proxy. At a reduction potential of −485 mV, metronidazole is reduced by Fd/Fld, inhibiting nitrogen fixation but not altering the respiration or ATP/ADP ratio (34–36). Therefore, an increased ratio of reduced:oxidized Fd/Fld intensifies the detrimental effect of metronidazole, decreasing growth rates (37). Strain Δ*rnf1* can grow on a higher concentration of metronidazole than WT and Δ*fix* strains (Figure 1). The growth pattern shows that the Δ*rnf1* strain has less reduced Fd/Fld compared to both WT and Δ*fix,* which suggests Rnf plays a dominant role in the electron transport mechanism, as there is a lower reduced:oxidized Fd/Fld ratio when Fix is the only enzyme present to generate low potential electrons (Δ*rnf1* strain).

**Figure 1.**
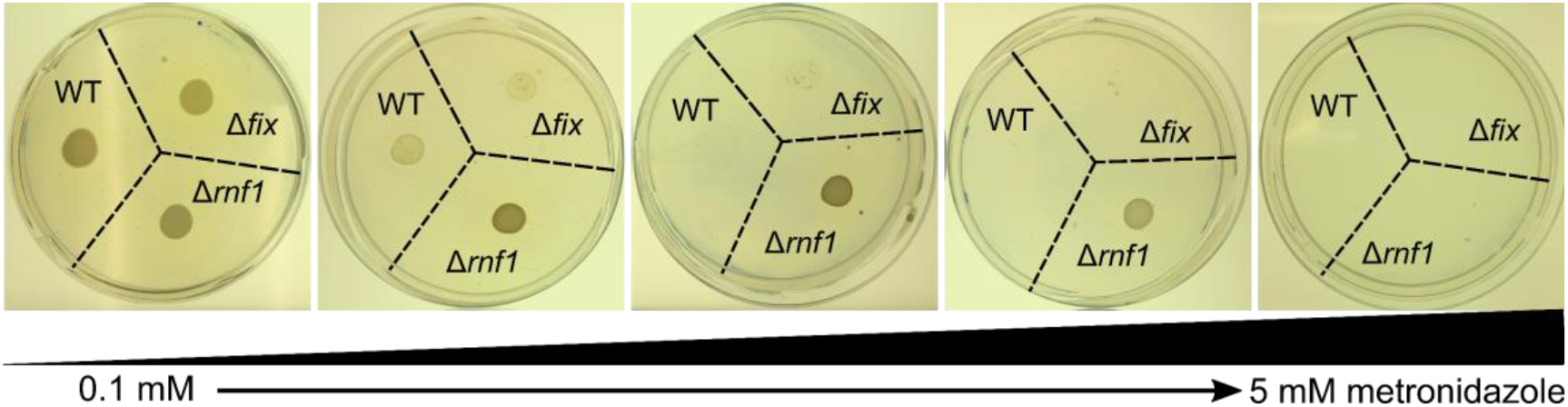
Metronidazole susceptibility assay of on agar plate of B medium with addition of 0.1, 0.5, 1, 2, and 5 mM of metronidazole. The Δ*rnf1* strain was able to grow on a concentration of 2 mM metronidazole where as WT and Δ*fix* strains were susceptible to metronidazole as low as 0.5 mM.

While redox homeostasis within the cell differs between strains, other energy forms might be compensating to maintain a standard growth rate. The ATP/ADP ratio was determined for cells growing in high aeration conditions. WT and Δ*fix* strains maintain an ATP/ADP ratio of 4-5, but the Δ*rnf1* strain has a significantly higher ATP/ADP ratio of ~6.5 (Figure 2) (Mann-Whitney, Wt vs. Rnf p = 0.0096).

**Figure 2.**
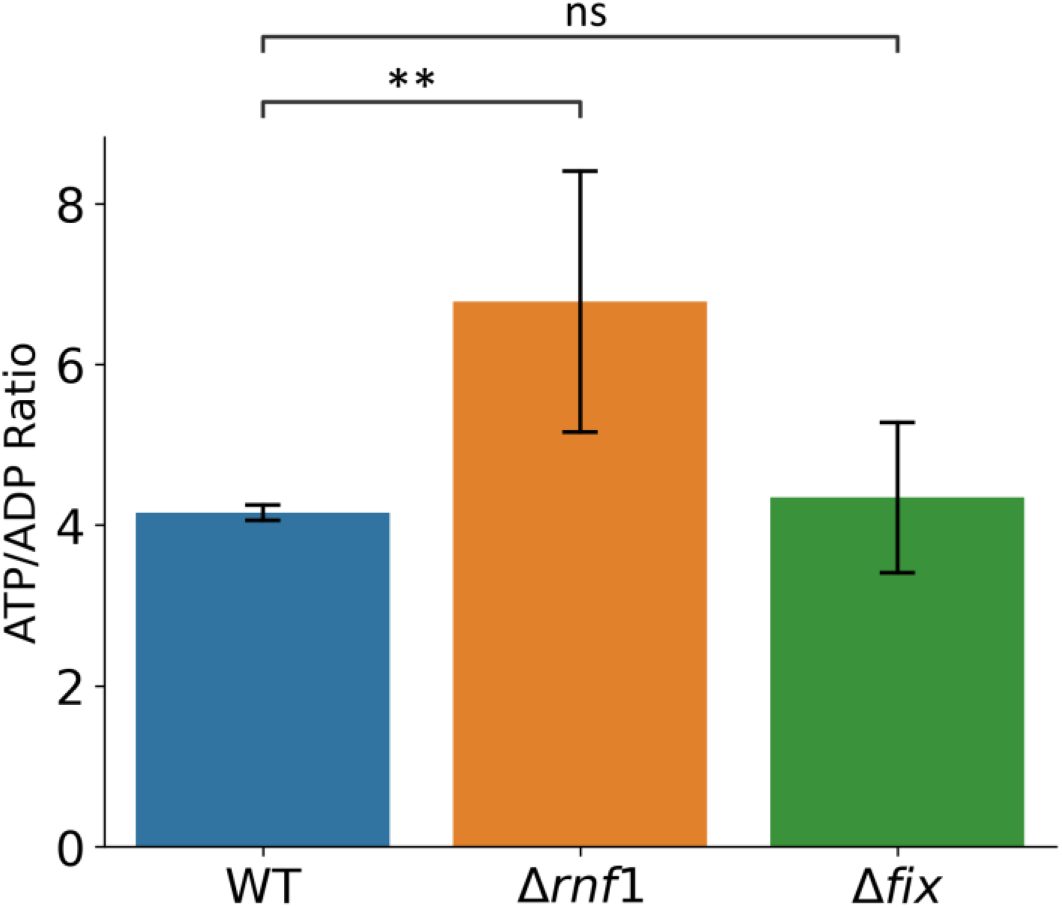
ATP/ADP ratio were measured by luciferase assay under high aeration conditions. WT and Δ*fix* cells have no significant difference in ATP/ADP ratio but Δ*rnf1* increase ATP/ADP ratio significantly compare to WT (Mann-Whitney, p = 9.9 × 10^−3^).

Increased ATP concentration could be from a rise in *pmf* and synthesis of ATP or a general decrease of ATP hydrolysis. ATP regeneration is highly correlated with nitrogenase activity, and energy supply has a more significant effect on nitrogenase activity than oxygen concentration (38, 39). The maintenance of *pmf* at a low enough potential is a significant factor in maximizing oxygen reduction; therefore, favoring the use of Rnf and its ability to consume *pmf*. In tandem with the evidence of the metronidazole experiment, we propose that Rnf is the primary mechanism for generating low potential electrons under standard laboratory conditions of excess oxygen and carbon.

While Rnf is preferred under oxygen-excess conditions and Fix is required for maximal growth under oxygen-limited conditions, the preferred electron transport mechanism to maximize energy production remains unknown. *A. vinelandii* has alternative nitrogenases used when Mo is limited within the environment (40). These alternative nitrogenases operate with different reaction stoichiometries compared to the reaction stoichiometry of Mo-nitrogenase and require more ATP and reductant to reduce N_2_ (41, 42). Flux balance analysis has shown that *A. vinelandii*, under alternative nitrogenase conditions, shifts electron flux from oxygen reduction to Fd/Fld and ATP production to meet the increased energy demand (31). Fix has been proposed to be the preferred mechanism in these conditions, as previous work showed that *fix* gene transcripts are up-regulated in alternative nitrogenase conditions (17). To test the effects of metal availability on electron transport strains, growth rates were determined in conditions lacking Mo but added V to express the V-nitrogenase and conditions without Mo or V to express the Fe-only nitrogenase. The overall energetic cost of nitrogen fixation rises 30% for V-nitrogenase and 60% for Fe-only nitrogenase, leading to slower growths and longer generation times (Table 2) (41, 42). The Δ*rnf1* strain grows slightly slower than the WT in these conditions. However, the Δ*fix* strain grows significantly slower when either V-nitrogenase or Fe-only nitrogenase is expressed. Slower growth of the Δ*fix* strain and up-regulation of *fixABCX* gene transcripts under alternative conditions imply Fix is the primary donor of electrons to nitrogenases in these conditions (17).

**Table 2.**
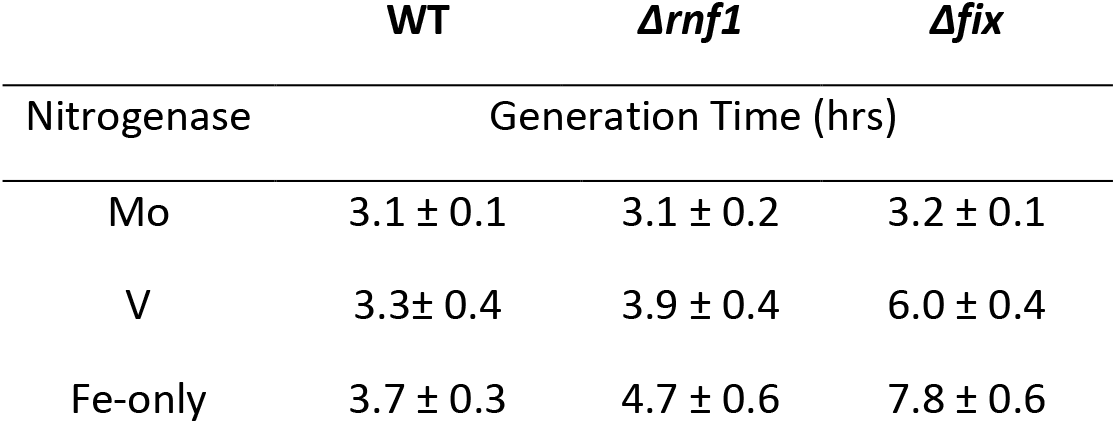
Generation times of WT, Δ*rnf1 and* Δ*fix* strains under Mo-dependent, V-dependent and Fe-only diazotrophic growth in high aeration conditions. Switching to V or Fe-only conditions increases the cost of nitrogen fixation. Cells were grown in 200 mL of media in a 500 mL baffled Erlenmeyer flask at 200 RPM. For Mo conditions standard B media was used, V conditions replaces Na_3_MoO_4_ with Na_3_VO_4_. Fe-only conditions have no Mo or V added.

|  | WT | *Δrnf1* | *Δfix* |
| --- | --- | --- | --- |
| Nitrogenase | Generation Time (hrs) |  |  |
| Mo | 3.1 ± 0.1 | 3.1 ± 0.2 | 3.2 ± 0.1 |
| V | 3.3 ± 0.4 | 3.9 ± 0.4 | 6.0 ± 0.4 |
| Fe-only | 3.7 ± 0.3 | 4.7 ± 0.6 | 7.8 ± 0.6 |

While experimental evidence shows that Rnf and Fix have different roles within the cell, both enzymes can be expressed simultaneously and interact with oxidative phosphorylation and the respiratory protection mechanisms that support nitrogen fixation (17, 24). To understand this more deeply, a simple thermodynamic model was constructed to determine the effects of *pmf* on the thermodynamic favorability of Rnf. First, the stoichiometry of protons translocated was calculated from previous physiological measurements to be ~5 H^+^/2e^−^ (43). This allows for the determination of ΔG of the Rnf reaction at a specific proton electrochemical gradient (Δμ_H+_). The Δμ_H+_ (KJ/mol) can be translated as the electromotive force of *pmf* (mV), giving a range of RNF’s thermodynamic feasibility. The model shows that a *pmf* of 79 mV is required for Rnf to be thermodynamically feasible, meaning Rnf should have a negative ΔG in almost all scenarios as ATP synthesis cannot be maintained at such a low *pmf* (44). While the ΔG of the Fix reaction was determined to be −36.7 kJ∙mol^−1^, the *pmf* should be near 145 mV for Rnf to be more favorable than Fix (Figure 3) (22). The stoichiometry of Rnf has only been measured for Na^+^ export in *Acetobacterium woodii* as 3 Na^+^/ 2e^−^ (45). While the pumping of Na^+^ ions to create Na^+^ motive force might not be comparable to the consumption of *pmf* in an aerobe, multiple stoichiometries were tested. Modeling different proton stoichiometries also shows that increased protons translocated (6 H^+^/ 2e^−^) would lower the *pmf* requirement to compete with Fix thermodynamically while fewer protons translocated (3-4 H^+^/ 2e^−^) would increase the *pmf* requirement.

**Figure 3.**
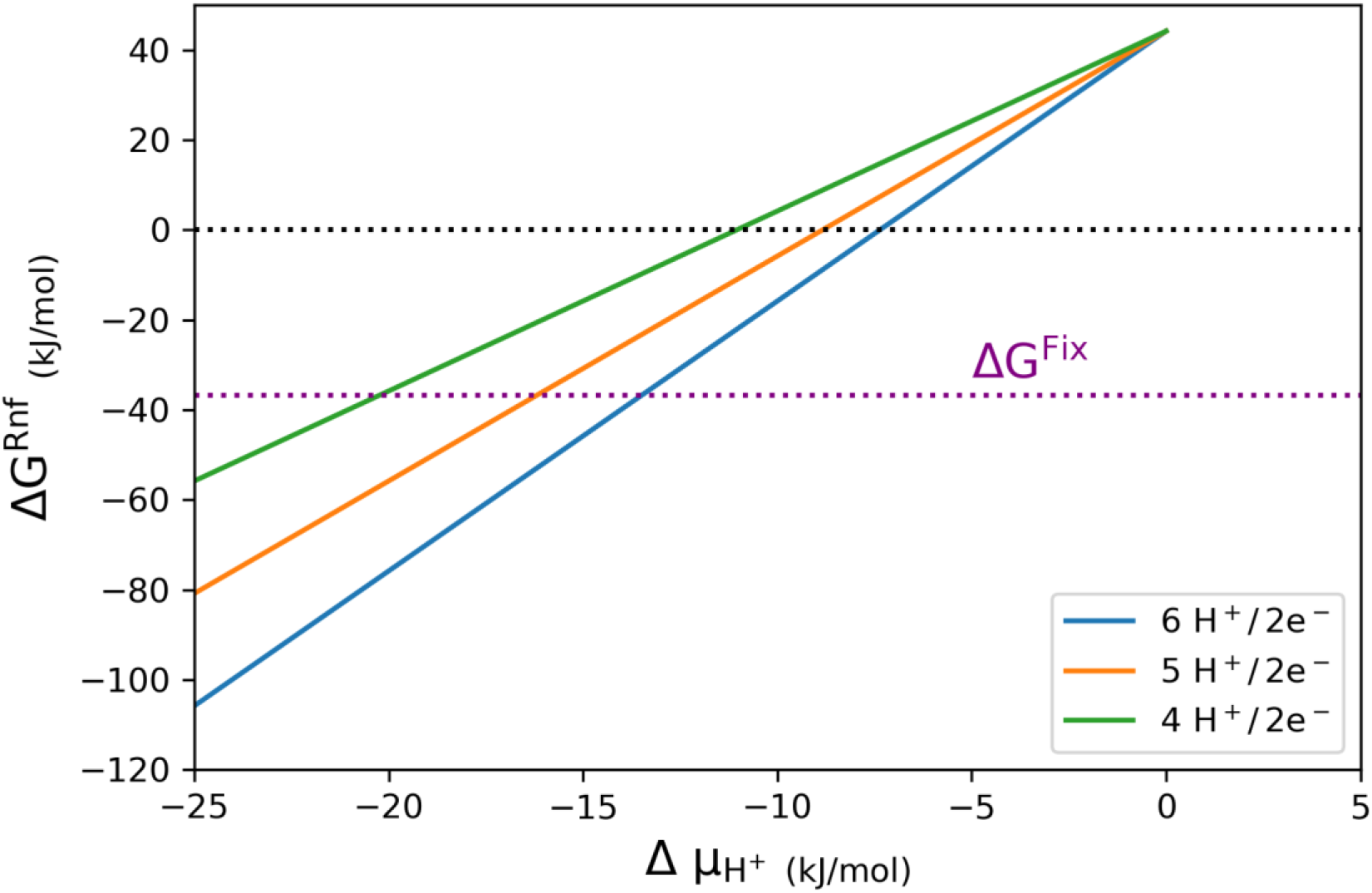
ΔG of Rnf as a function of *pmf.* The ΔG of Rnf is dependent on the magnitude of the *pmf.* The ΔG of Rnf (KJ/mol) was compared to the proton electrochemical gradient (Δμ_H+_, KJ/mol) at different proton translocation stoichiometries (colors). As a comparison the ΔG of Fix is labeled at −36.7 KJ/mol as calculated by (22).

The *pmf* for *A. vinelandii* under oxygen-limited conditions has not been quantified, but other studies measuring *pmf* in gram-positive bacteria show no change from high to low oxygen concentrations (46, 47). This maintenance of *pmf* between oxygen concentrations is compensated by adjusting the Δψ and ΔpH components to maintain ATP synthesis. Since Rnf has been biochemically characterized as a Na^+^-transporter in other bacteria, it may be affected by the Δψ component of *pmf* in *A. vinelandii* (48–50). While the nitrogen status transcriptionally regulates Rnf1, it is also post-translationally regulated by the glutamine:2-oxoglutarate ratio. In *Azoarcus sp BH72,* Rnf activity is inhibited in high nitrogen or anaerobiosis shock by regulatory protein GlnK (51). A similar mechanism in *A. vinelandii* could affect Rnf efficiency under low oxygen or low *pmf* conditions.

A final model can be proposed for the relationship of Fix and Rnf in the context of aerobic nitrogen fixation (Figure 4). Fix is favored as the mechanism for electron transport to nitrogenase under oxygen-limited conditions or when extra energy is required for nitrogen reduction. The bifurcation of electrons to quinone supplements ATP production while still reducing Fd/Fld; this maximizes the energy production needed for nitrogen fixation. While under standard laboratory conditions of substrate excess, maximizing *pmf* consumption plays a role in maintaining respiratory protection. Here, we see that Rnf is favored as its mechanism consumes *pmf* and may help balance ATP production and oxygen consumption. While each enzyme can sustain electron transport to nitrogenase, the corresponding mechanism of action defines that enzyme’s role. This system of intrinsic regulation through each electron transport mechanism may allow for fast balancing of the electron transport system to maintain efficient nitrogen fixation.

**Figure 4.**
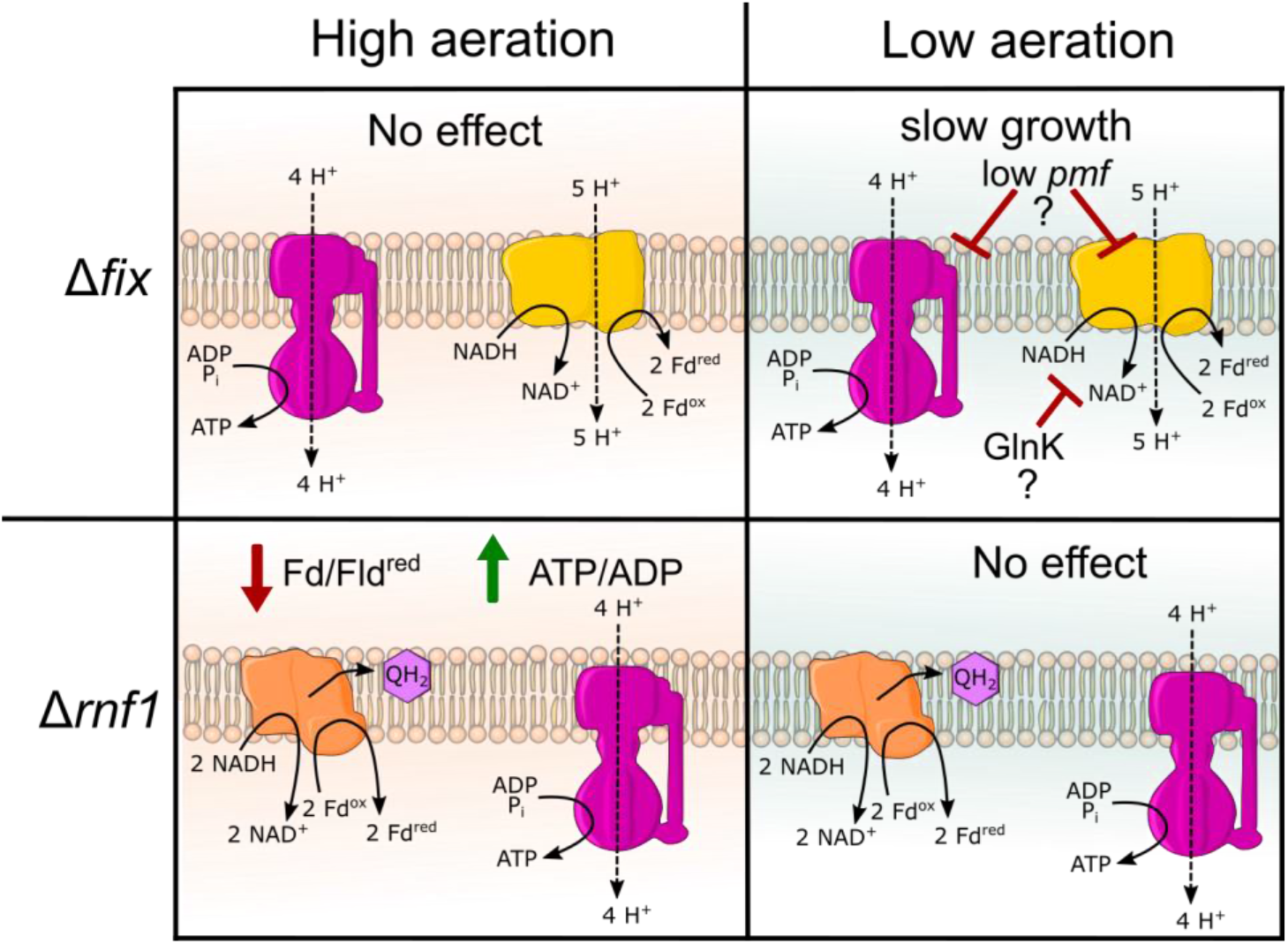
Overall roles of Rnf and Fix under different aeration conditions. The two component system of Rnf (yellow) and Fix (orange) are simultaneously expressed during nitrogen fixation. The deletion of either enzyme does not cause growth effects under standard laboratory conditions of high aeration. The Δ*rnf1* strain does impact the overall redox homeostasis and increases ATP through ATP synthase (purple) under high aeration. Under low aeration the Δ*fix* strain grows significantly more slowly, as Rnf is the only way to produce reduced Fd/Fld. Thermodynamic modeling suggests that this may be because of a lowering of the *pmf* or may be interacting with nitrogen regulator protein GlnK.

## Materials and Methods

Strains and growth conditions- Wild-type *A. vinelandii* strain DJ was used (52), Δ*rnf1* and Δ*fix* strains were obtained from previous work (16, 22). For the standard growth assays, cells were grown aerobically at 30°C in Burk’s sucrose medium (B medium) or Burk’s sucrose medium supplemented with 10 mM ammonium acetate (BN medium) (53). Cells were grown in 500-mL baffled Erlenmeyer flasks with 200-mL liquid medium. Flasks were incubated on a rotary shaker at 200 RPM for high aeration or 90 RPM for low aeration. Growth was measured by optical density at 600 nm in a Thermo scientific tabletop spectrophotometer. To avoid the known effects of *rnf1* deletions on nitrogenase derepression, all strains were first transferred to B-media agar plates then inoculated into B-media as a starter culture. This adaptation to nitrogen-fixing conditions eliminated the lag in growth previously seen in Δ*rnf1* strains (16).

The alternative nitrogenases were expressed following the methods from Sippel *et al.* (54). Briefly, DMSO stocks (Mo-containing) were grown to the mid-log phase, streaked onto agar plates made from B-V media, containing 0.01 mM Na_3_VO_4_ instead of Na_3_MoO_4_. Next, colonies grown on B-V plates were used to inoculate ammonia-supplemented vanadium media (BN-V). This process was repeated five times until RT-PCR of *nifK* and *vnfK* genes determined only vanadium nitrogenase expression. Next, the protocol was repeated to express the Fe-only nitrogenase, but with media lacking both Mo and V. All glassware was acid washed, and plastics were soaked in a 1 mmol/L EDTA bath. Once alternative nitrogenase expression was confirmed, DMSO stocks were made for future use.

To confirm the oxygen concentration phenotype, cell growth of WT, Δrnf1, and Δfix was tested under the same shaking rate but in different atmospheres. Strains were grown at 30°C in 2 mL of B medium in six-well plates shaken at 500 RPM in a double orbital pattern. The optical density at 600 nm was taken every 30 minutes using a BMG Clariostar plate reader (BMG Lab Tech, Ortenberg, Germany). In addition, an atmosphere control unit kept a constant 19.4 % O_2_ (atmospheric) or 10 % O_2_ by N_2_ dilution.

Physiological assays – Enzylight ADP/ATP ratio kit (BioAssay Systems, Hayward, CA, USA) was used to measure ATP/ADP ratio by following the manufacturer’s protocol and measuring luminescence on a Clariostar plate reader (BMG Lab Tech, Ortenberg, Germany). Metronidazole sensitivity was used as a proxy for redox potential within the cell (55). B medium plates were made as described above but with the addition of 0.1, 0.5, 1, 2, and 5 mM metronidazole. Cells were grown to an OD of 0.5 in standard B medium, and 25 μL of cells were placed onto the plates. Cell growth was determined after 48 hrs.

Thermodynamic model – The stoichiometry of Rnf in *A. vinelandii* is unknown and must be determined to investigate the effects of *pmf* on nitrogen-fixing conditions.

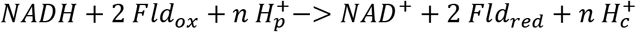

Two electrons from NADH are transferred to two molecules, Fd/Fld, with the movement of protons from the periplasm 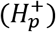 to the cytoplasm 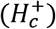. The actual redox potential of the NAD^+^/NADH redox couple was calculated from the *in vivo* ratios of 12.5 (measured in Garcia et al. (26)) and the Nernst equation,

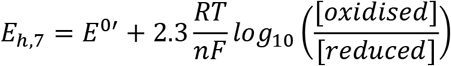

where the *E*_*h*,7_ is the actual redox potential, *R* is the gas constant, *F* is the Faraday constant, *T* is the temperature in °K, and *n* is the number of electrons transferred. *E*_*h*,7_ was calculated to be −287 mV when *E*^0′^, the standard redox potential, was −320 mV.

Next, the difference in the actual redox couples, Δ*E*_*h*_, was calculated by the difference between NAD^+^/NADH midpoint potential (determined above) and the Fld midpoint potential of −483 mV (56) to give a value of −196 mV.

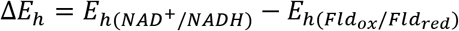

The Gibbs energy change (Δ*G*) accompanying electron transfer between couples can be calculated at 37.82 kJ∙mol^−1^.

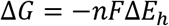

The *pmf* was measured in *A. vinelandii* by Laane et al. (43) as the electric potential (Δψ) at 106 mV and the change in pH (Δ*pH*) at 0.45 pH. The proton electrochemical gradient 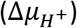 in kJ∙mol^−1^ can be calculated from Δψ and ΔpH in order to translate the *pmf* into electron millivolts (mV).

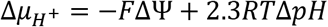

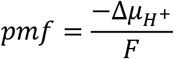

Finally, to calculate the protons translocated for a thermodynamically favorable reaction, both the 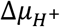 and Δ*p* can be used in different equations to produce 4.96 H^+^/2e^−^ or 4.91 H^+^/2e^−^, respectively. Where the protons translocated (n) is calculated by the energy difference (KJ/mol) or voltage (mV)

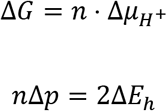

Thus giving the thermodynamically calculated stoichiometry for Rnf:

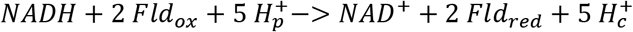

Since a single *pmf* value was used to determine this stoichiometry, a plot was created to see *pmf’s* effect on the ΔG of the Rnf reaction. In addition, multiple stoichiometries were plotted to ensure a biologically relevant range of thermodynamic feasibility. All equations were from Bioenergetics 4 (57).

## Acknowledgments

This work was supported by the US DOE, Office of Science, Office of Basic Energy Sciences under award DE-SC0018143 to J. W. P. Partial salary support for J. W. P. was supported by the United States Department of Agriculture National Institute of Food and Agriculture, Hatch umbrella project #1015621.

---

**Supplementary Table 1.** Generation times of WT Δ*rnf1 and* Δ*fix* strains grown in 2 mL of B medium in a six-well plate. Plates were shaken at 300 RPM in a double orbital rotation and atmosphere was controlled with added nitrogen.

|  | WT | *Δrnf1* | *Δfix* |
| --- | --- | --- | --- |
|  | Generation Time (hrs) |  |  |
| 19.4 % O<sub>2</sub> (atm) | 3.1 ± 0.0 | 3.0 ± 0.1 | 4.0 ± 0.0 |
| 10% O<sub>2</sub> | 4.9 ± 0.1 | 4.5 ± 0.1 | 8.7 ± 0.2 |

